# Finding analytic stationary solutions to the chemical master equation by gluing state spaces at one or two states recursively

**DOI:** 10.1101/113340

**Authors:** X. Flora Meng, Ania-Ariadna Baetica, Vipul Singhad, Richard M. Murray

## Abstract

Noise is often indispensable to key cellular activities, such as gene expression, necessitating the use of stochastic models to capture its dynamics. The chemical master equation (CME) is a commonly used stochastic model that describes how the probability distribution of a chemically reacting system varies with time. Knowing analytic solutions to the CME can have benefits, such as expediting simulations of multiscale biochemical reaction networks and aiding the design of distributional responses. However, analytic solutions are rarely known. A recent method of computing analytic stationary solutions relies on gluing simple state spaces together recursively at one or two states. We explore the capabilities of this method and introduce algorithms to derive analytic stationary solutions to the CME. We first formally characterise state spaces that can be constructed by performing single-state gluing of paths, cycles or both sequentially. We then study stochastic biochemical reaction networks that consist of reversible, elementary reactions with two-dimensional state spaces. We also discuss extending the method to infinite state spaces and designing stationary distributions that satisfy user-specified constraints. Finally, we illustrate the aforementioned ideas using examples that include two interconnected transcriptional components and chemical reactions with two-dimensional state spaces.

**Subject Areas:** Systems biology, synthetic biology, biomathematics, bioengineering

## 1 Introduction

Stochastic fluctuations in the levels of cellular components are essential to biological processes such as gene expression coordination and cellular probabilistic differentiation [1,2]. There are two commonly used approaches to modelling the time evolution of a spatially homogeneous mixture of molecular species that interact through a set of known chemical reactions [3]. The deterministic formulation specifies the time-rates-of-change of the molecular concentrations of component species with a set of coupled differential equations and assumes continuous variations in the molecular concentrations [4]. In contrast, the stochastic formulation considers the time evolution of the probability distribution of molecular compositions which is modelled by a set of coupled linear differential equations [5, 6]. Although the deterministic approach is adequate in many cases, its assumption that a chemical reaction system evolves deterministically and continuously can be invalid at low molecular counts due to experimental evidence of stochastic effects and integer-valued molecular counts [7]. In such situations, stochastic modelling becomes necessary [8-10].

The chemical master equation (CME) is a stochastic model that describes how the probability distribution of molecular counts of chemical species in a reacting system varies as a function of time [7, 11]. The CME describes a continuous-time Markov jump process: each state represents the molecular counts of all component species, and transitions between states correspond to changes in molecular counts via chemical reactions [12]. Analytic solutions to the CME are important in biological engineering for a number of reasons. For instance, simulations of biochemical reaction networks that are multiscale in time can be expedited by incorporating analytic solutions of the fast time-scale dynamics [13]. Analytic stationary solutions also enable accurate analysis and design of the effect of parameter values on the stationary behaviour of a reaction system [13]. In general, analytic forms of transient and stationary solutions to the CME for arbitrary reaction network topologies are still unknown, mainly because the number of states increases exponentially with the number of component species [14]. However, there are known results for some simple biochemical reaction networks with particular initial conditions [14], such as mass-conserving [15] linear reaction systems with multinomial initial distributions [16, 17], and linear reaction systems with Poisson initial distributions [18]. In the absence of analytic solutions, Monte Carlo algorithms, such as Gillespie’s stochastic simulation algorithm (SSA) [3, 5], are used to approximate the solution to the CME. Nevertheless, the computational cost of SSA becomes enormous when there are numerous reactions present in the system, and the method does not guarantee error bounds on the approximate solution [19]. Mélykúti et al. [13,20] recently proposed a technique for determining the analytic stationary distributions of the CME for stochastic biochemical reaction networks whose state spaces can be constructed by gluing two finite, irreducible state spaces at one or two states sequentially. The stationary distribution on the combined state space is a linear combination of the equilibria of the single Markov jump processes [13]. An analogous method for gluing state spaces at two states was developed in [20]. Mélykúti et al.’s gluing technique forms a basis for the construction of recursive algorithms that provides a fast way to compute analytical expressions for stationary distributions on large state spaces. To illustrate this recursion graphically, we can imagine gluing three triangles together at their vertices to form a triangular grid, and by forming increasingly larger triangular grids using existing ones, we can rapidly construct very large grids.

The gluing technique has a number of potential advantages over existing methods. The simplest approach that one might take to compute an analytic stationary distribution of a continuous-time Markov jump process is to solve for a left null vector of the transition rate matrix [12]. However, the dimension of the matrix is almost always infinite, often making the calculation exceedingly difficult. There are numerical methods such as the finite state projection (FSP) algorithm [19] that can be used to truncate infinite state spaces. However, it is unclear how to employ the FSP algorithm to obtain analytic stationary solutions. The gluing technique can provide analytic solutions for finite state spaces, and may be the basis for recursive algorithms that derive analytic solutions for infinite state spaces. The gluing technique also provides a link between stationary distributions to the CME and the graphical structure of the state space. Furthermore, predicting the dynamic behaviour of a biological network from that of its constituent modules is a central yet generally unsolved problem in systems and synthetic biology [21-24]. The development of the gluing technique can potentially contribute to computing analytic solutions using its recursive property [20].

In this work, we explore the capabilities of Mélykúti et al.’s gluing technique and introduce recursive algorithms that use the technique to compute the stationary solution to the CME. We first introduce basic notions, including the CME model, the relationship between Markov jump processes and graph theory, and Mélykúti et al.’s gluing technique. We then use graph theory to formally characterise the set of state spaces that can be constructed by carrying out single-state gluing of paths, cycles or both sequentially. We subsequently demonstrate the single and double-state gluing techniques with simple chemical reaction systems. In addition, we discuss extending the method to infinite state spaces and designing stationary distributions that satisfy user-specified constraints. Finally, we illustrate the aforementioned ideas using examples that include two interconnected transcriptional components [25].

## 2 Background

To introduce Mélykúti et al.’s [13,20] gluing technique, we first present the CME and then explain its connection with graph theory.

### 2.1 The chemical master equation

The chemical master equation describes the time evolution of a chemically reacting system as a continuous-time Markovian random walk in the space of molecular counts of chemical species [5, 11]. Consider a system with *n* chemical species 𝒮_1_,…, 𝒮_n_ in a well-stirred solution of fixed volume and temperature. We denote the molecular count of each species 𝒮_*i*_ by *X_i_* ∈ ℤ_≥0_. Suppose that the molecular counts of these chemical species are specified by the initial state *X*^0^ ∈ ℤ_≥0_^*n*^ at the time *t*_0_ ≥ 0 and they change to other states *X* = (*X*_1_, …, *X_n_*) via *m* possible chemical reactions ℛ_1_,…, ℛ_*m*_. For each reaction ℛ_*j*_, let *a_j_* (*X*) be the associated *propensity function* and *v_j_* ∈ ℤ ^*n*^ the *state-change vector* [18], which we will define as follows. At state *X*, the probability that a single occurrence of ℛ_*j*_ takes place in an infinitesimal interval *dt* is given by *a_j_*(*X*)*dt*, and *v*_*j*_ is the vector of induced changes in *X*. Therefore, the probability that a particular reaction ℛ_*j*_ occurs first, i.e. the random walk jumps to the state *X* + *v*_*j*_ next, is equal to 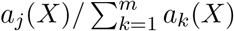. We assume *mass-action* kinetics in this work. The reaction equations of the zeroth, first, and second-order reactions have left-hand side of the forms 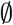, 𝒮_*i*_, 𝒮_*i*_ + 𝒮_*l*_ (*l* ≠ *i*), and 2𝒮*_i_*. These reactions have propensity functions *κ*, *κX*_*i*_, *κX_i_X_l_*, and 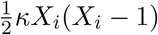, respectively, with an appropriate rate constant *κ* > 0.

Let Pr(*X, t* | *X*^0^, *t*_0_) denote the probability of the Markov jump being at state *X* at time *t* > *t*_0_, conditional on the initial state *X*^0^. The CME is then a set of coupled linear differential equations, with the equation for state *X* being

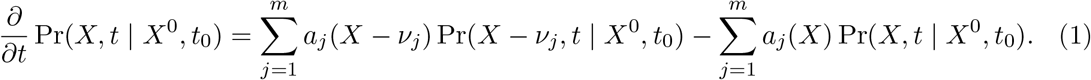

The first summation term on the right-hand side of equation (1) corresponds to reactions via which the random walk can jump to state *X* in one step, and the second summation term considers reactions from which the random walk can leave *X* in one step [14].

### 2.2 The relationship between continuous-time Markov jump processes and graph theory

In this subsection, we explain the connection between graph theory and the Markov jump process that underlies the CME. We only consider reversible chemical reactions. The state space of the Markov jump process described by the CME naturally gives an undirected graph without self-loops (i.e. edges that connect a vertex to itself): vertices represent states, and an edge exists between a pair of vertices if and only if there exists a reversible reaction that allows transition between the two corresponding states.

For instance, gene expression regulation can be described as the reversible binding of transcription factors with copies of a gene. Let 𝒯, 𝒢, and 𝒢* denote the transcription factor, the free gene, and the gene-transcription factor complex, with molecular counts *T*, *G*, and *G\**, respectively. Let *κ_b_* and *κ_d_* be the binding and unbinding rate constants. The reaction equation is then given by

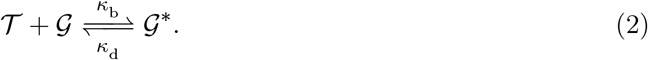

If we assume the reaction system to be closed (i.e. mass-conserving), then mass conservation leads to the relation *T* + *G* + 2*G\** = *C*, where *C* ∈ ℤ_≥ 0_ is a constant that is determined by the initial condition. For instance, if the initial condition is (*T, G, G\**) = (1, 1, 2), then the state space with the corresponding transition rates is and the corresponding graph is a path of length 3.

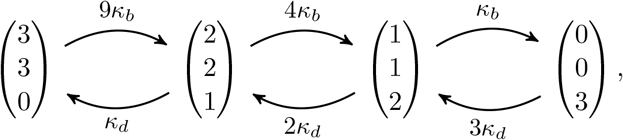

In the rest of this work, we refer to the graph that is given by a state space with corresponding transition rates simply as the state space for brevity. In Section 2.3, we discuss deriving the stationary distribution of a continuous-time Markov jump process by gluing two state spaces together at one or two vertices.

### 2.3 The stationary distribution of a continuous-time Markov jump process glued together from two state spaces at one or two vertices

Having introduced the CME and its connection with graph theory, we are ready to present the technique proposed by Mélykúti et al. [13,20].

Mélykúti, Hespanha, and Khammash developed a technique in [13] for solving the stationary distribution of a continuous-time Markov jump process by gluing the state spaces of two finite, irreducible, continuous-time Markov jump processes at exactly one vertex. Mélykúti, and Pfaffelhuber [20] considered gluing state spaces at two vertices simultaneously. In both papers, vertices are glued together if and only if they correspond to the same state. Without introducing any new jumps or losing any existing jumps, a new Markov process is naturally identified on the combined state space. No existing literature has studied gluing at more than two vertices.

We first consider the one-vertex gluing technique in [13]. Suppose that A and B are two continuous-time Markov jump processes with finite, irreducible state spaces. Suppose that process A has a known stationary distribution *ξ*^*A*^ on states indexed by {1, 2,…, *r*}, and process B has a known stationary distribution *ξ*^*B*^ on states indexed by {1, 2,…, *s*}. The order in which the states are indexed does not affect the result. Without loss of generality, we assume gluing the two state spaces at state r of process *A* and state 1 of process *B*. We relabel the states by keeping the indices of process *A* the same and increasing the indices of process *B* by *r* – 1. The new Markov process has a unique stationary distribution [13] given by

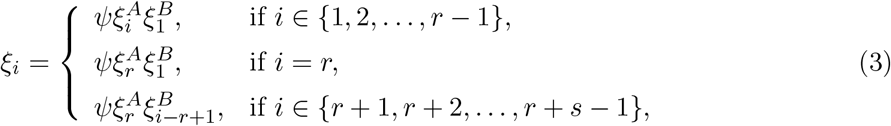

where 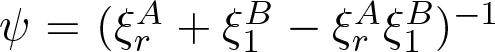 is a normalising constant. Figure 1 illustrates the one-vertex gluing technique using the example in Section 2.2.

**Figure 1:**
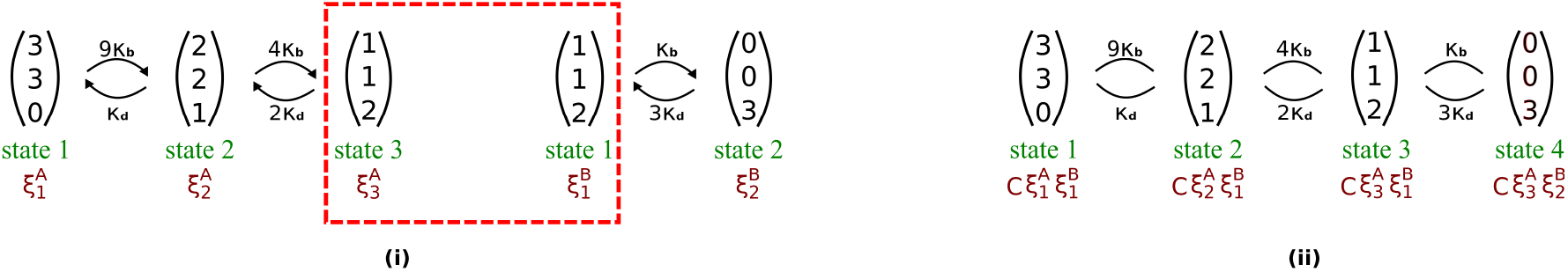
An illustration of the one-vertex gluing technique in [13]. (i) The state spaces of two Markov jump processes *A* and *B* with known stationary distributions (*ξ*_1_^*A*^, *ξ*_2_^*A*^, *ξ*_3_^*A*^) and (*ξ*_1_^*B*^, *ξ*_2_^*B*^), respectively. State 3 of *A* and state 1 of *B* represent the same state, and all the other states are distinct, so we glue the two state spaces at these two states. (ii) The resulting state space and the corresponding stationary distribution, where 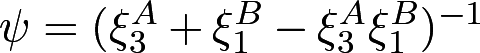 is a normalising constant.

When the two processes to be glued together have exactly two pairs of identical states, we employ the two-vertex gluing introduced in [20]. With the same setting as before, we now glue state *r* – 1 of process A to state 1 of process *B*, and state *r* of process *A* to state 2 of process *B*. We relabel the states by keeping the indices of process A the same and increasing the indices of process B by *r*–2. The glued vertices are not necessarily consecutive in index, but we assume the index introduced for simplicity of notation. In general, deriving the stationary distribution of the new Markov process obtained by two-vertex gluing entails lengthy calculations. However, [20] proves that if the proportionality condition specified by the equation

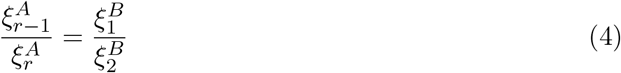

is satisfied, then the stationary distribution on the combined state space is

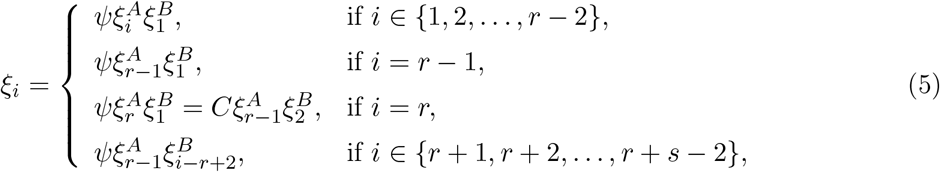

where 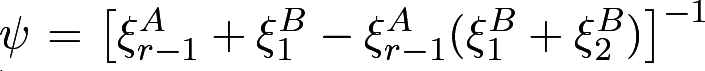 is a normalising constant. The condition in Equation (4) is necessary for the stationary distribution on the combined state space to preserve the proportions of probability within each Markov jump process. It is interesting that this condition is also sufficient.

## 3 Characterising graphs that can be obtained by gluing paths, cycles, or both, at one vertex sequentially

In this section, we first introduce definitions in graph theory based on [26], and then use them to characterise the set of state spaces that can be obtained by gluing paths, cycles, or both, at one vertex sequentially. This set of state spaces provides a natural starting point for studying Mélykúti et al.’s gluing technique [13,20] for two main reasons. First, analytic solutions are already known for the stationary distributions on path-like and circular state spaces [12, 27, 28]. Second, one-vertex gluing involves only simple arithmetic and is computationally efficient. We find that (i) graphs obtained by gluing paths at one vertex sequentially are trees, (ii) graphs obtained by gluing cycles at one vertex sequentially are “trees of cycles”, and (iii) graphs obtained by gluing paths and cycles at one vertex sequentially are “trees of trees and cycles”. We give formal propositions describing these results in this section and proofs in the electronic supplementary material.

Let *X*^(*k*)^ = {*A* ⊆ *X* : |*A*| = *k*} be the set of *k*-element subsets of *X*. A *graph G* is defined by an ordered pair (*V*, *E*) where *V*(*G*) ≠ 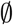 is a finite set, called the *vertex set*, and *E*(*G*) ⊆ *V*^(2)^ is an *edge set*. We consider undirected graphs without self-loops in this work, as explained in Section 2.2. In other words, if {*u*, *υ*} ∈ *E*(*G*), then *u* ≠ *υ* and we can denote {*u*, *υ*} by *uυ* or *υu* equivalently. If *uυ* ∈ *E*(*G*), then vertices *u* and *υ* are *adjacent* and are both *endvertices* of edge *uυ*. A *path of length n*, called an *n-path*, is a graph with the vertex set {*υ_i_* : *i* = 1, 2,…, *n* + 1} and the edge set {*υ_i_ υ_i_* + _1_ : *i* = 1, 2, …,*n*}. By convention, a 0-path is a vertex. A *cycle of length n*, called an *n-cycle*, is a graph with the vertex set {*υ_i_* : *i* = 1, 2,…, *n*} and the edge set {*υ_i_υ_i_*_+1_ : *i* = 1, 2,…, *n* − 1} ∪ {*υ_n_υ_1_*}. A graph is *connected* if any two distinct vertices are joined by a path. A connected and acyclic graph is called a *tree.* Furthermore, a graph *H* is a *subgraph* of *G*, written as *H* ⊆ *G*, if *V*(*H*) ⊆ *V*(*G*) and *E*(*H*) ⊆ *E*(*G*). Suppose that *G* is a connected graph. For our purposes, we define the graph *G* − *H* as the graph obtained by first deleting edges of *H* from *G* and then removing isolated vertices of the remaining graph (see

Figure 2 for an example). Similarly, if *G* has three or more vertices and *e* ∈ *E*(*G*), then the graph *G* – *e* is obtained by first deleting the edge *e* from *G* and then removing any isolated endvertex of *e* in the remaining graph.

**Figure 2:**
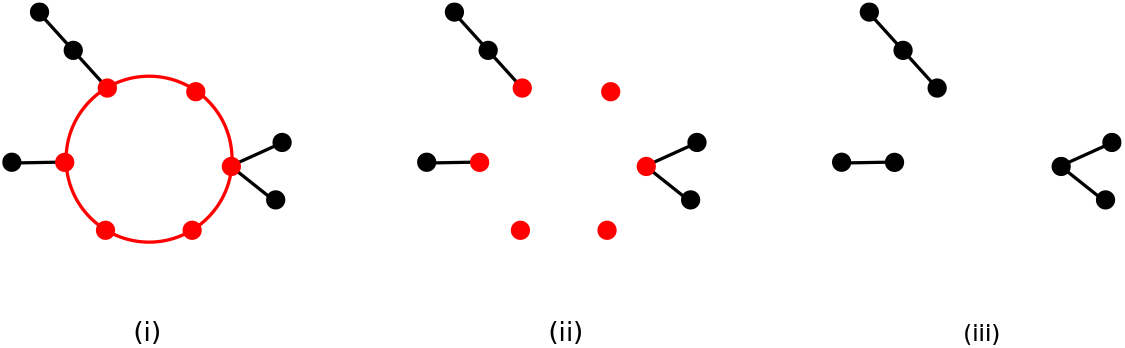
An illustration of removing a subgraph from a graph. (i) An example of a graph G with a circular subgraph *H* (in red). To obtain *G* – *H*, (ii) delete the edges of *H* from *G*, and (iii) remove isolated vertices from the remaining graph.

### Proposition 1.

*A graph can be obtained by gluing paths together at one vertex sequentially if and only if the graph is a tree.*

Figure 3 gives an example of a tree and one way to construct the graph by gluing paths at one vertex sequentially.

**Figure 3:**
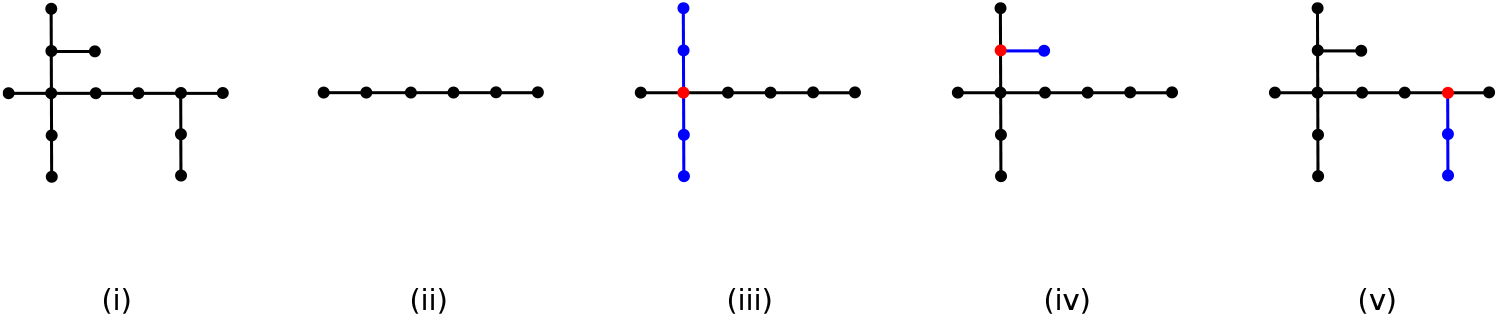
(i) An example of a tree. (ii)-(v) One way to construct the tree by gluing paths together at one vertex sequentially. New components are in blue. Graphs are glued together at red vertices.

The *degree* of a vertex *υ* ∈ *V*(*G*), denoted by *d_G_*(*υ*), is the number of distinct vertices that are adjacent to *υ* in graph *G*. A vertex with zero degree is said to be *isolated.*

### Proposition 2.

*A graph can be obtained by gluing cycles together at one vertex sequentially if and only if the graph satisfies all of the following conditions:*

i. *the graph is connected,*
ii. *every vertex has an even degree, and*
iii. *any two distinct cycles have at most one common vertex.*

A graph that satisfies all of the conditions in Proposition 2 can be thought of colloquially as a “tree of cycles”. Figure 4 gives an example of a tree of cycles and one way to construct the graph by gluing cycles at one vertex sequentially. For a formal definition, see the electronic supplementary material.

**Figure 4:**
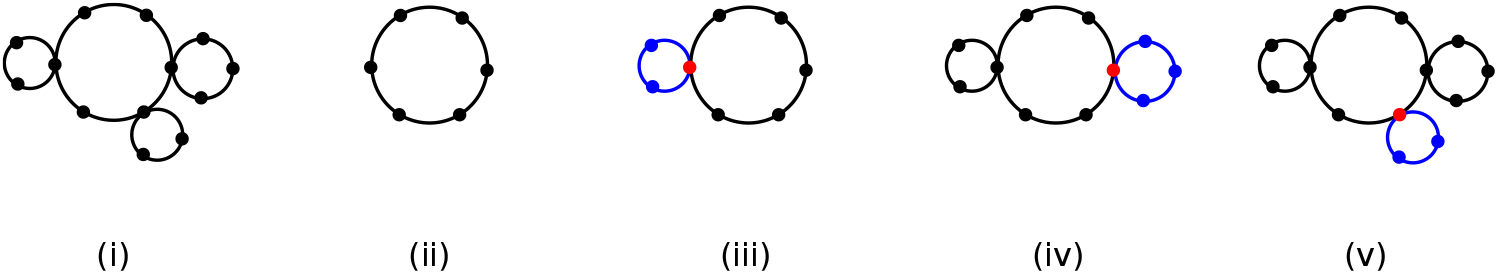
(i) An example of a tree of cycles. (ii)-(v) One way to construct the graph by gluing cycles together at one vertex sequentially.

### Proposition 3.

*A graph can be obtained by gluing paths and cycles together at one vertex sequentially if and only if the graph satisfies both of the following conditions:*

i. *the graph is connected, and*
ii. *any two distinct cycles share at most one common vertex.*

Intuitively, a graph that satisfies both of the conditions in Proposition 3 is a “tree of trees and cycles” (see Figure 5 for an example).

**Figure 5:**
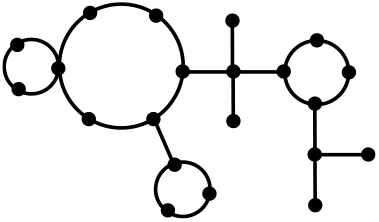
An example of a tree of trees and cycles.

The one-vertex gluing technique [13] provides an easy method for computing the stationary distributions of biochemical reaction systems whose state spaces are characterised by Propositions 1-3. We note that any state space can be constructed by gluing simple graphs together at one or two vertices consecutively, such as by adding one edge at a time. However, this strategy entails a large number of steps to build large graphs and thus is computationally inefficient. Moreover, deriving the stationary distribution of the new Markov process obtained by two-vertex gluing is complicated in general, except when the proportionality condition in equation (4) holds. In Section 4, we explore the development of recursive algorithms based on the one and two-vertex gluing techniques in [13] and [20] through a set of biochemical reaction systems.

## 4 Computing analytic stationary solutions to the CME using recursive algorithms

In this section, we propose recursive algorithms that apply Mélykúti et al.’s gluing technique to find stationary solutions to the CME of a set of biochemical reaction systems. In Section 4.1, we define the set of chemical reactions with finite state spaces for which our algorithms are intended and demonstrate with examples. In Section 4.2, we derive the analytic stationary solution of two interconnected transcriptional components, for which the state space is infinite, and discuss designing stationary distributions with desired properties.

### 4.1 Closed systems with reversible, elementary reactions

In this subsection, we shall focus on a set of chemical reactions that have finite state spaces, which can be constructed by using Mélykúti et al.’s gluing technique finitely many times. We study a biochemical reaction system with an infinite state space in Section 4.2.

We first define an *elementary reaction* in Definition 1 for the purposes of our study.

#### Definition 1.

Consider three distinct chemical species 𝒮_1_, 𝒮_2_, and 𝒮_3_. For *i* = 1, 2, and 3, let *α*_*i*_, *β*_*i*_ ∈ {0, 1, 2}. A reaction of the form

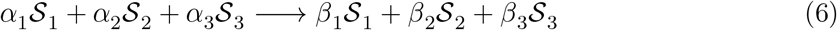

is an *elementary reaction* if 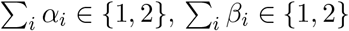, and 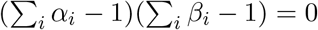.

The reactions specified by Definition 1 can be categorised into five types: 𝒮_1_ → 𝒮_2_, 𝒮_1_ → 2𝒮_2_, 𝒮_1_ → 𝒮_2_ + 𝒮_3_, 2𝒮_2_ → 𝒮_1_, and 𝒮_2_ + 𝒮_3_ → 𝒮_1_. These reactions can be seen as building blocks for more complex reactions. For instance, 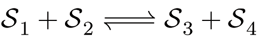 is an approximation to the reactions 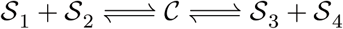, where C is an intermediate.

We study reversible, elementary reactions in closed systems in this subsection. The state spaces of these biochemical reaction systems are determined by the initial state and the number of reversible reactions present.

#### 4.1.1 One reversible reaction

There are three kinds of closed systems with exactly one reversible, elementary reaction, which we list in Table 1. The state spaces of these reactions are paths, the lengths of which are determined by the initial state. We first present the known results in [27] about the stationary distributions on finite, path-like state spaces. We then propose a recursive algorithm that uses the one-vertex gluing technique.

**Table 1:** Systems with exactly one reversible, elementary chemical reaction.

|  |
| --- |
| $\mathcal{S}_1 \rightleftharpoons \mathcal{S}_2$ |
| $2\mathcal{S}_1 \rightleftharpoons \mathcal{S}_2$ |
| $\mathcal{S}_1 + \mathcal{S}_2 \rightleftharpoons \mathcal{S}_3$ |

Let *N* ∈ ℤ _>0_ and consider a continuous-time Markov jump process on a path-like state space with vertices indexed by {0, 1,…, *N*}. Suppose that *p_i_* is the transition rate from *i* to *i* + 1 for *i* ∈ {0, 1, …, *N* − 1}, and *q_i_* from *i* to *i* – 1 for *i* ∈ {1, 2, …, *N*}. The stationary distribution (*ξ*_0_, *ξ*_1_,…, *ξ*_*N*_) is given by the relation 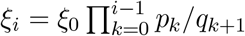 and the normalisation equation 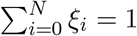.

For *x* ≥ 0, let ⌊*x*⌋ denote the integer part of *x*. Alternatively, the stationary distribution can be obtained by gluing paths of lengths 1, 2, 4, …, 2^⌊log_2_*N*⌋−1^ at one endvertex sequentially and then adding a path of length *N* − 2^⌊log_2_*N*⌋^. The stationary distribution on a path of length *N* − 2^⌊log_2_*N*⌋^ can be obtained using a similar strategy. Consider the example in Section 2.2 with initial state (*T, G, G\**) = (7, 7, 0). Figure 6 illustrates graphically our recursive algorithm to obtain the stationary distribution on the state space, which is a path of length 7.

**Figure 6:**
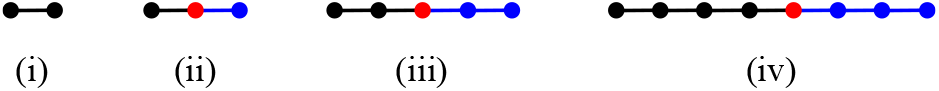
A graphical demonstration of a recursive algorithm that uses the one-vertex gluing technique in [13] to solve the stationary distribution on a path-like state space. New components are in blue. Graphs are glued together at red vertices.

#### 4.1.2 Two reversible reactions

There are eight types of closed systems with exactly two reversible, elementary reactions, which we list in Table 2. The state spaces of these reactions are diagonally truncated grids, the sizes of which depend on the initial state. We propose a recursive algorithm that applies Mélykúti et al.’s gluing technique to compute analytic stationary distributions on such state spaces.

**Table 2:** Systems with exactly two reversible, elementary chemical reactions.

|  |
| --- |
| $\mathcal{S}_1 \rightleftharpoons \mathcal{S}_2 \rightleftharpoons \mathcal{S}_3$ |
| $2\mathcal{S}_1 \rightleftharpoons \mathcal{S}_2 \rightleftharpoons \mathcal{S}_3$ |
| $\mathcal{S}_1 \rightleftharpoons 2\mathcal{S}_2 \rightleftharpoons \mathcal{S}_3$ |
| $2\mathcal{S}_1 \rightleftharpoons \mathcal{S}_2 \rightleftharpoons 2\mathcal{S}_3$ |
| $\mathcal{S}_1 + \mathcal{S}_2 \rightleftharpoons \mathcal{S}_3 \rightleftharpoons \mathcal{S}_4$ |
| $\mathcal{S}_1 \rightleftharpoons \mathcal{S}_2 + \mathcal{S}_3 \rightleftharpoons \mathcal{S}_4$ |
| $\mathcal{S}_1 + \mathcal{S}_2 \rightleftharpoons \mathcal{S}_3 \rightleftharpoons \mathcal{S}_4 + \mathcal{S}_5$ |
| $\mathcal{S}_1 + \mathcal{S}_2 \rightleftharpoons \mathcal{S}_3 \rightleftharpoons 2\mathcal{S}_4$ |

Consider a closed system with two coupled monomolecular reversible reactions

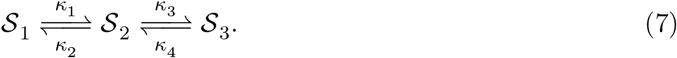

The total number of molecules, denoted by *N*, is a constant that is determined by the initial state. It is sufficient to denote each state by a 2-vector (*X*_1_, *X*_2_) as *X*_3_ can be calculated using *X*_3_ = *N* − *X*_1_ − *X*_2_. Figure 7 shows the state space when *N* = 3 and the corresponding graphical representation. All closed systems with exactly two reversible, elementary reactions have state spaces of such a diagonally truncated grid structure.

**Figure 7:**
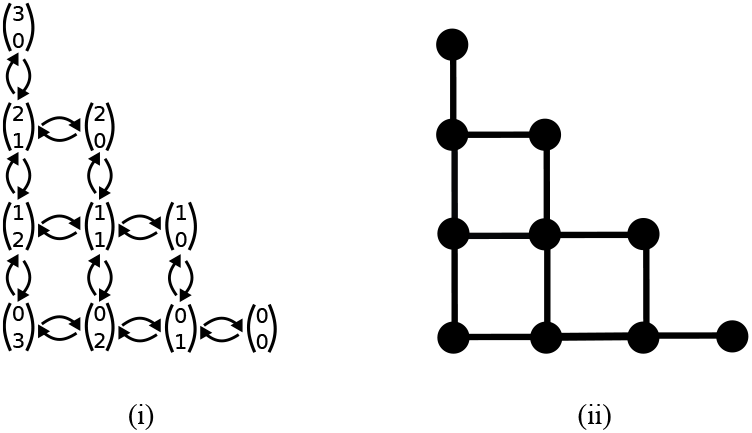
(i) The state space of a closed system with two coupled monomolecular reversible reactions. Formation and degradation of species 𝒮_1_ correspond to moving up and down the state space, respectively. Formation and degradation of species 𝒮_3_ correspond to moving right and left, respectively. (ii) The corresponding graphical representation.

When *N* = 1, the state space is simply a 3-path. For integer *N* ≥ 2, we propose an algorithm that takes *N* and *κ*_*j*_, where *j* ∈ {1, 2, 3, 4}, as inputs and returns the stationary distribution of the corresponding chemical reaction system. The algorithm constructs the state space recursively by sequentially gluing together small graph components. Figure 8 demonstrates the case when *N* = 3. Specifically, we construct the state space by first gluing together three L-shaped components sequentially at one vertex and then checking the proportionality condition, as given in equation (4), for the remaining missing edges. The method is applicable to the general case for any integer *N* ≥ 2. We label states in the state space from left to right and from bottom to top so that the index increases naturally as the graph grows.

**Figure 8:**
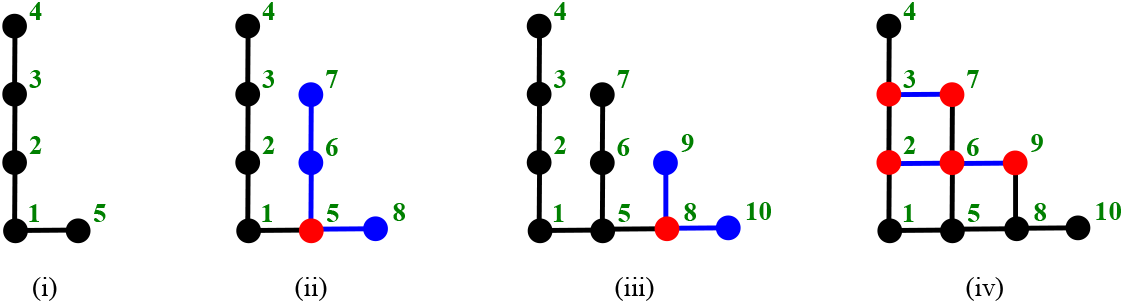
A graphical demonstration of constructing the state space of a closed system with two coupled monomolecular reversible reactions when *N* = 3. Vertices are labelled in green. New components are in blue. Graphs are glued together at red vertices. In Subfigures (ii) and(iii),we glue together three L-shaped components sequentially at one vertex. In Subfigure (iv), blue edges are added one at a time.

In general, we can construct the state space by first gluing together *N* L-shaped components sequentially at one vertex and then checking the proportionality condition, as given in equation (4), for all missing edges. For each L-component, we call the state with the lowest index state 1, the state with the second lowest index state 2, and so on. For *x*_2_ ∈ {1, 2, 3, …, *N*}, the stationary distribution *ξ*^*x*_2_^ on an L-shaped component with state 1 being (*X*_1_, *X*_2_) = (0, *x*_2_) is given by

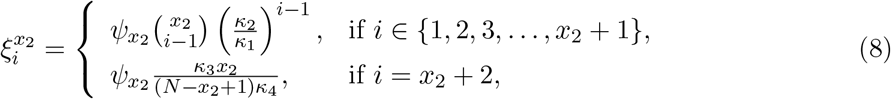

where *ψ*_*X*_2__ is a normalising constant. After gluing together *N* of these L-shaped components sequentially at one vertex, the graph contains gill of the states from the state space. In order to complete the construction, we need to add 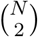 edges by gluing at the two endvertices of each missing edge simultaneously. The proportionality condition, as given in equation (4), holds at all missing edges (i.e. all edges between states (*i* − 1, *x*_2_ − *i* + 1) and (*i* − 1, *x*_2_ − *i*) for *x*_2_ ∈ {2, 3, 4, …, *N*} and *i* ∈ {2, 3, 4,…, *x*_2_}). Therefore, the stationary distribution of the orignal state space is the same as that of the state space constructed with L-shaped components which is given by the equation

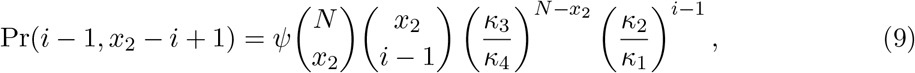

with the normalising constant 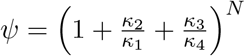 for *x*_2_ ∈ {1, 2, 3,…, *N*} and *i* ∈ {1, 2, 3,…, *x*_2_ + 1}. Moreover, the state with the highest index has probability of 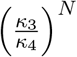 in the stationary distribution.

#### 4.1.3 Three reversible reactions

A *planar graph* is a graph that can be drawn on the plane in such a way that edges intersect only at their endvertices [26]. There are three possible closed systems with exactly three reversible, elementary reactions, the state spaces of which are planar graphs. We list these reaction systems in Table 3. In such a system, each elementary reaction gives rise to the same chemical change as the overall effect of the other two reactions. If this condition does not hold or a chemical reaction system contains more than three reversible, elementary reactions, then its state space is not a planar graph, and our recursive algorithm does not apply.

**Table 3:** Systems with exactly three reversible, elementary chemical reactions.

|  |  |  |
| --- | --- | --- |
| $\mathcal{S}_1 \rightleftharpoons \mathcal{S}_2$ | $\mathcal{S}_2 \rightleftharpoons \mathcal{S}_3$ | $\mathcal{S}_3 \rightleftharpoons \mathcal{S}_1$ |
| $\mathcal{S}_1 \rightleftharpoons 2\mathcal{S}_2$ | $2\mathcal{S}_2 \rightleftharpoons \mathcal{S}_3$ | $\mathcal{S}_3 \rightleftharpoons \mathcal{S}_1$ |
| $\mathcal{S}_1 \rightleftharpoons \mathcal{S}_2 + \mathcal{S}_3$ | $\mathcal{S}_2 + \mathcal{S}_3 \rightleftharpoons \mathcal{S}_4$ | $\mathcal{S}_4 \rightleftharpoons \mathcal{S}_1$ |

Consider a closed system with three coupled monomolecular reversible reactions

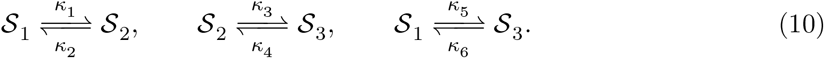

Figure 9 shows the state space when *N* = 3 and the corresponding graphical representation. All closed systems with exactly three reversible, elementary reactions as given in Table 3 have state spaces of such a diagonally truncated grid structure.

**Figure 9.**
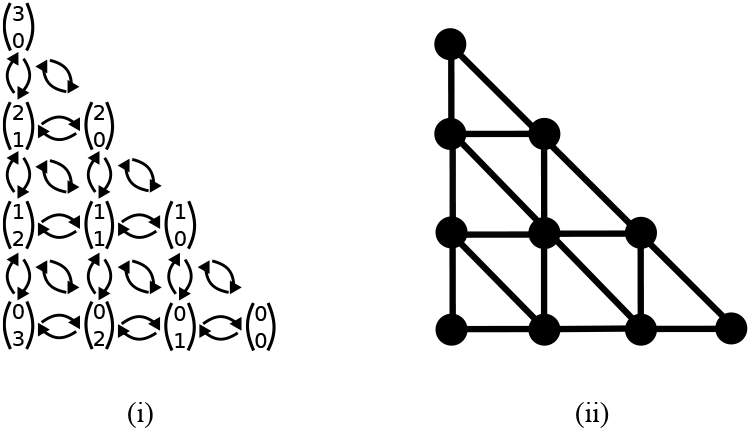
(i) The state space of a closed system with three coupled monomolecular reversible reactions as given in Equation (10). In addition to the transitions illustrated in Figure 7, diagonal moves denote conversion between species 𝒮_1_ and 𝒮_3_. (ii) The corresponding graphical representation.

We observe that the recursive algorithm introduced in Section 4.1.2 contructs part of the state space of the system with reactions given in Equation (10). If the proportionality condition in Equation (4) is also satisfied for all of the diagonal edges, then the three-reaction system has the same stationary distribution as that of the two-reaction system. Therefore, a closed system with three coupled monomolecular reversible reactions as given in Equation (10) has the stationary distribution that is specified by Equation (9) if the reaction rate constants are compatible in the sense that

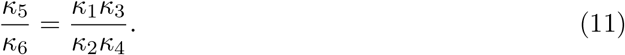

If Equation (11) does not hold, then it is an open question whether the chemical reaction system has a stationary distribution or not.

### 4.2 Stationary distributions of two interconnected transcriptional components

In this subsection, we study the stationary behaviour of two interconnected transcriptional components [25] by developing a recursive algorithm that is analogous to the algorithms introduced in Section 4.1. Specifically, we first model the cascade of two connected transcriptional components using the CME. We then apply the gluing technique on a finite subset of the infinite state space, providing an error bound on the approximate solution. We subsequently obtain the analytic stationary solution of the CME by taking the size of the subset to infinity, arriving at the same results as in [25]. Finally, we discuss designing stationary distributions that satisfy user-specified constraints by searching over the parameter space of the biochemical reactions.

#### 4.2.1 Modelling two interconnected transcriptional components using the CME

We first present the chemical reactions that are introduced in [25] to model two interconnected transcriptional components and then provide the corresponding CME model.

Figure 10 illustrates a transcriptional component connected to a downstream component. The upstream component comprises the constitutive expression of a transcription factor *Ƶ*, which subsequently binds reversibly to a promoter 𝒫 at the downstream component. Let 𝐶 denote the complex formed by *Ƶ*, and 𝒫. The reactions are then given by the chemical equations

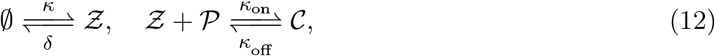

where *κ* > 0, *δ* > 0, *κ*_on_ > 0, and *κ*_off_ > 0 are the corresponding reaction rates. Let *P, C*, and *Z* be the numbers of 𝒫, 𝐶, and *Ƶ*, respectively. Since the total amount of DNA is conserved, it always holds that *P* + *C* = *N* for some constant *N* that is determined by the initial condition.

**Figure 10:**
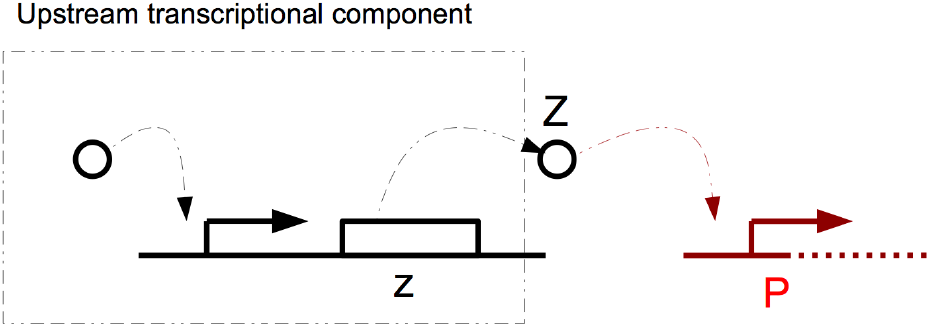
Two interconnected transcriptional components. The figure is reproduced from [25] with permission.

If the initial condition is (*C, Z, P*) = (*c_0_, z_0_, p_0_*) ∈ ℤ_≥0_^3^, then the conservation constant is given by *N* = *c*_0_ + *p*_0_. It is sufficient to denote each state by a 2-vector (*C*, *Z*) as *P* can be calculated using *P* = *N* − *C*. Since there can be arbitrarily many transcription factors, the reaction system described by equation (12) has an infinite state space, as illustrated in Figure 11.

**Figure 11:**
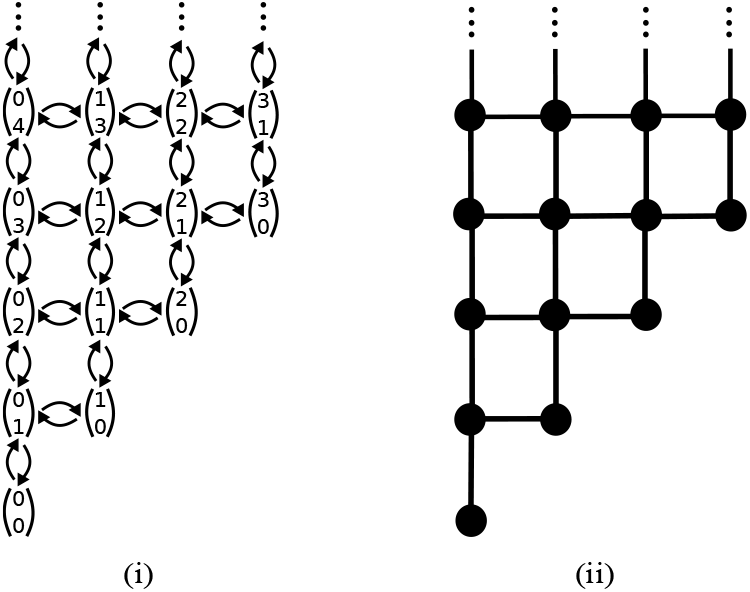
(i) The infinite state space of two interconnected transcriptional components with reactions given by Equation (12) when *N* = 3. Expression and degradation of transcription factors, *Ƶ*, correspond to moving up and down the state space, respectively. Formation and decomposition of complexes, 𝒞, correspond to moving right and left, respectively. (ii) The graphical representation of the state space given in (i).

For simplicity, we assume that the chemical reaction system has volume *V* = 1. Let Pr(*c*, *z*) be the probability that the stochastic process is in state (*c*, *z*) ∈ ℤ_≥0_^2^, at time *t* > 0, conditional on the initial state (*C, Z*) = (*c*_0_, *z*_0_). Each state (*c, z*) contributes exactly one linear ordinary differential equation to the CME, which is given by

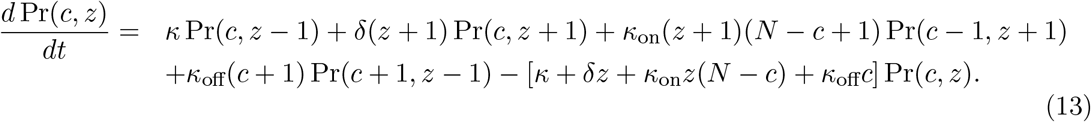

The CME of two interconnected transcriptional components comprises infinitely many differential equations that have the form of Equation (13). Therefore, we need to truncate the infinite state space to a finite subset that can then be recursively constructed by applying Mélykúti et al.’s gluing technique finitely many times.

#### 4.2.2 Truncating infinite state spaces with guaranteed accuracy

Applying the initialisation step of Munsky and Khammash’s finite state projection (FSP) algorithm [19], we approximate the stationary distribution on the infinite state space. Specifically, depending on the desired accuracy of the approximate solution, we truncate the infinite state space by imposing a maximum value, *M* ∈ ℤ_>0_, on the number of transcription factors considered. For instance, Figure 12 presents the truncated state space of Figure 11 when *M* = 2.

**Figure 12:**
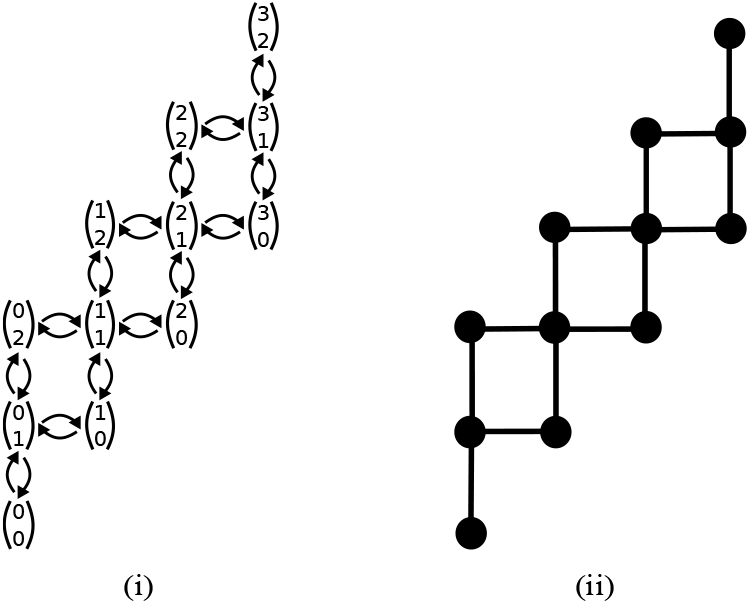
(i) The truncated state space of two interconnected transcriptional components when *N* = 3 and *M* = 2. (ii) The graphical representation of the truncated state space given in (i).

We now derive *M* as a function of the desired accuracy. Let Ω be the infinite state space, and suppose that *ϵ* ∈ [0, 1] is the level of acceptable error. We choose *M* such that the total probability that the finite state approximation fails to capture is at most *ϵ*. Let Ω_f_ ⊂ Ω denote the corresponding truncated finite state space. In other words, the total probability accounted by Ω_f_^c^ ≡ Ω\Ω_f_ is at most ϵ. In order to calculate the total probability that is accounted for by the states in Ω_f_^c^ at equilibrium, we regard the states in Ω_f_ as one state, called *ω*_f_, and states in Ω_f_^c^ as state *ω*_c_. The transition rate from *ω*_f_ to *ω*_c_ is the sum of the transition rates from Ω_f_ to Ω_f_^c^, and vice versa. Therefore, the transition rate from *ω*_f_ to *ω*_c_ is *κ*(*N* + 1), and the reverse transition rate is *δ*(*M* + 1)(*N* + 1). At equilibrium, the probabilities of *ω*_f_ and *ω*_c_ are 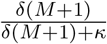 and 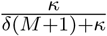, respectively. We choose *M* ∈ ℤ_>0_ such that 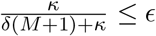. Hence, we choose

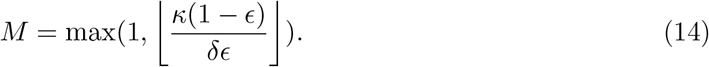

#### 4.2.3 Analytic stationary solutions on infinite state spaces

We propose a recursive algorithm that, together with asymptotic analysis, yields the analytic stationary solution to the CME of two interconnected transcriptional components. The algorithm takes *N*, *M*, *κ*, *δ*, *κ*_on_, and *κ*_off_ as inputs and returns the stationary distribution on the corresponding truncated finite state space. The algorithm constructs the truncated state space recursively by gluing small graph components together sequentially. In the limit of *M* → ∞, the stationary distribution of the truncated state space converges to that of the original infinite state space.

We demonstrate the recursive algorithm for the case when *N* = 3 and *M* = 2 in Figure 13, but the method is applicable to the general case of *N* ∈ ℤ_>0_ and *M* ∈ ℤ _>0_. Specifically, we construct the truncated state space by first gluing together three T-shaped components and a path sequentially at one vertex and then checking the proportionality condition, as given in equation (4), for the remaining missing edges. Similar to Section 4.1.2, we label states in the truncated state space from left to right and from bottom to top so that the index increases naturally as the graph grows.

**Figure 13:**
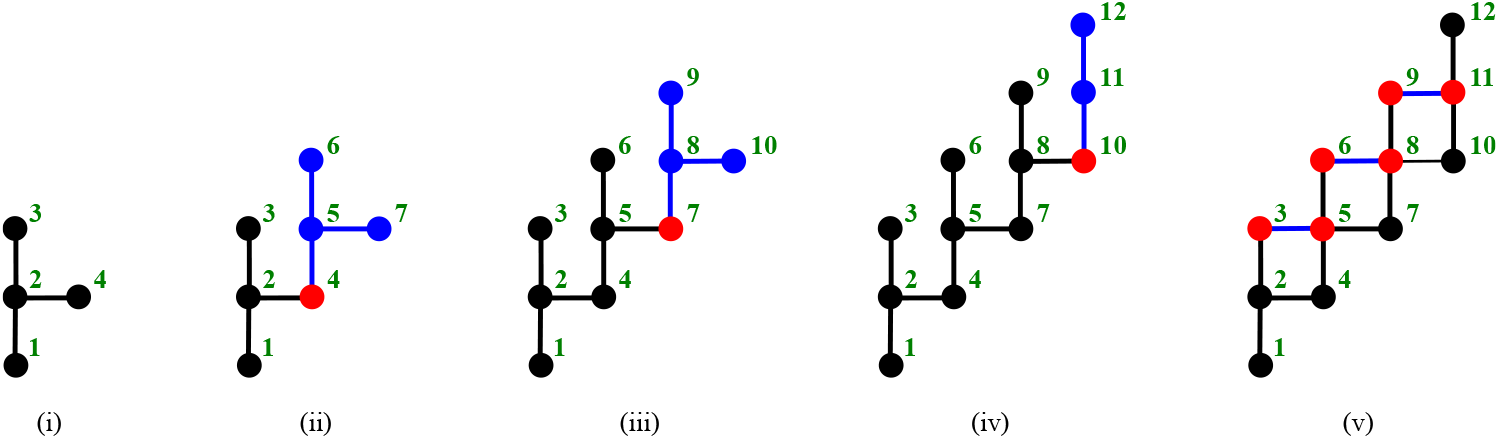
A graphical demonstration of constructing a truncated state space of two interconnected transcriptional components when *N* = 3 and *M* = 2. Vertices are labelled in green. New components are in blue. Graphs are glued together at red vertices. In Subfigures (i)-(iii), we glue together three T-shaped components sequentially at one vertex. In Subfigure (iv), we add a path in order to complete the vertex set. In Subfigure (v), blue edges are added one at a time.

In general, a T-shaped component consists of *M* + 2 states. There are three steps for constructing a truncated state space: (i) glue together *N* of T-shaped components sequentially at one vertex, (ii) add a path of length *M* to the existing state space at the vertex with the highest index, and (iii) check the proportionality condition, as given in equation (4), for all missing edges.

For *c* ∈ {0, 1, 2, …, *N* − 1}, the stationary distribution *ξ*^*c*^ on a T-shaped component for which state 1 (i.e. the vertex with the lowest index) represents (*C*, *Z*) = (*c*, 0) is given by

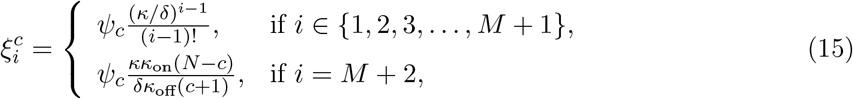

where *ψ_c_* is a normalising constant. The stationary distribution on the *M*-path is given by

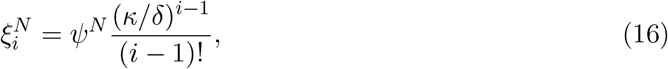

for *i* ∈ {1, 2, 3,…, *M* + 1}, where *ψ^N^* is a normalising constant.

After step (ii), the graph contains all the states from the truncated state space. In order to complete the construction, we need to add *N*(*M* − 1) edges by gluing at the two endvertices of each edge simultaneously. The proportionality condition, as given in equation (4), holds at all missing edges (i.e. all edges between states (*c*, *z*) and (*c* + 1, *z* − 1) for *c* ∈ {0, 1, 2, …, N−1} and *z* ∈ {2, 3, 4, …, *M*}). Therefore, the stationary distribution on the truncated state space is already obtained in step (ii). For instance, when *N* = 3 and *M* = 2, the stationary distributions of Subfigures 13(iv) and 13(v) are the same.

We may truncate the infinite state space to a finite subset by bounding the maximum value of *Z* as lim_*z* → ∞_ Pr(*c*, *z*) = 0 for *c* ∈ {0, 1, 2, …, *N*}. Moreover, the dependence of Pr(*c*, *z*) on *M* is only through the normalising constant. Therefore, by normalising over *z* ∈ ℤ_≥0_ and *c* ∈ {0, 1, 2, …, *N*}, we obtain the accurate stationary distribution of the original infinite state space, which is given by the equation

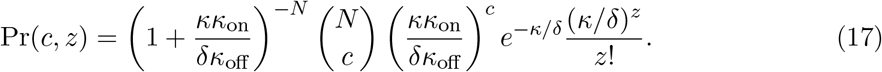

Equation (17) can be written as a product of a function of *c* and a function of *z*, which implies independence between the stationary behaviour of the upstream and downstream transcriptional components. In addition, the stationary distribution of random variable *C* is binomially distributed with the number of trials and success probability in each trial being *N* and [(*δκ*_off_)/(*κκ*_on_) + 1]^− 1^, respectively. The stationary distribution of random variable *Z* follows a Poisson distribution with mean *κ*/*δ*. Our results confirm theorem 5.1 in [25]. However, the proof of [25] relies on the deficiency zero theorem [29] and the theorem of product-form stationary distributions (theorem 4.1 of [30]).

#### 4.2.4 Designing stationary distributions using analytic solutions

Mathematical methods for designing distributional properties of stochastic biochemical systems have been developed in conjunction with the field of synthetic biology as a means to engineer predictable biochemical responses [31,32]. The stationary distribution design in this subsection is simpler than the general methods introduced in [31,32] because of the analytic solutions that we obtained using Mélykúti et al.’s gluing technique [13,20].

Computing stationary distributions in analytic form enables us to design the equilibrium behaviour of biochemical reaction networks simply by tuning their reaction rate parameters. For the two-component transcriptional system, this corresponds to altering the dilution rate or the degradation rate, making more RNA polymerases available to increase the transcription rate, or even choosing a transcription factor with the desired strength of binding and unbinding to DNA.

In this subsection, we focus on designing features of the stationary distribution of the two-component transcriptional system by adjusting the parameters of the biochemical reactions given in equation (12). We formulate and find solutions for two potential design problems in Propositions 4, 5, and Corollary 1. We consider two sets of design features, namely, the mean and the variance of the marginal probability distributions, and the location of the peak of the joint stationary distribution. For the latter, we prove that the stationary distribution has a unique global maximum if and only if the conditions in Proposition 5 are satisfied. Considering only stationary distributions with a unique global maximum simplifies the subsequent design of the peak's location. We present the proofs of Propositions 4 and 5 in the electronic supplementary material.

##### Proposition 4.

*Consider the system of two interconnected transcriptional components that are modelled by reactions as given in* *Equation* (12), *where κ* > 0, *δ* > 0, *κ_on_* > 0, *and κ*_*off*_ > 0 *are the corresponding reaction rate constants. Let P, Z, and C be the numbers of promoters 𝒫, transcription factors Ƶ, and 𝒫-Ƶ complexes 𝒞, respectively. Let* 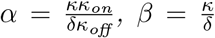, *and* 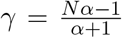, *where N is a constant given by N* = *P* + *C due to the conservation of DNA. In (i)–(iii), we set up and solve three design problems using the marginal stationary distributions of Z and C.*

i. *Since the marginal stationary distribution of Z is Poisson distributed, its mean and variance are equal. The design problem of fixing the mean of Z at an objective value μ_z_* > 0 *is feasible, and the solution is β = μ_z_, with N and the reaction rate constants being arbitrary otherwise.*
ii. *The design problem of setting the mean of C at an objective value μ_c_ ∈* (0, *N*) *is feasible, and the solution is 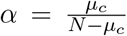 with N and the reaction rate constants being arbitrary otherwise.*
iii. *The design problem of choosing the variance of C to be an objective value σ_c_^2^* > 0 *is feasible if and only if* 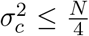, *and the solutions are* 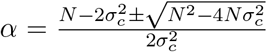, *with N and the reaction rate constants being arbitrary otherwise.*

We now consider designing the location of the peak of the joint stationary distribution as given in Equation (17). In Proposition 5, we find necessary and sufficient conditions for the existence of a unique global maximum and provide its location. We include the proof of Proposition 5 in the electronic supplementary material. Corollary 1 shows that the uniqueness of the peak simplifies the design of its location.

##### Proposition 5.

*Consider the system of two interconnected transcriptional components that are modelled by the reactions in* *Equation* (12). *With the same notation as in Proposition 4, the stationary distribution in* *Equation* (17) *has a unique global maximum if and only if N* > 1, *β* > 1, 0 < *γ < N* − 1, *and β, γ* ∉ *ℤ. In this case, the maximum is at* (*c*, z\**) *=* (⌊*γ*⌋ + 1, ⌊*β*⌋).

##### Corollary 1.

*Under the constraints N > 1, β >* 1, 0 < *γ* < *N* − 1, *and β, γ* ∉ *ℤ, designing the location of the unique global maximum of the two-component transcriptional system modelled by the reactions in* *Equation* (12) *is equivalent to finding N, β, and γ such that* (⌊*γ*⌋ + 1, ⌊*β,*⌋) *is the objective location.*

When the global maximum of the two-component transcriptional system exists and is unique, there are infinitely many parameter values that can lead to the objective location of the peak, and no closed-form solutions exist because of the floor-function form. Since any a α, β, γ, and *N* that satisfy the conditions in Corollary 1 would lead to the desired location, this implies the relative insensitivity of the peak’s location with respect to experimental inaccuracies.

Propositions 4, 5, and Corollary 1 illustrate that analytic solutions can greatly facilitate the design of stationary distributions. Design problems can be formulated as feasibility problems over decision variables, which are a α, β, γ, and *N* in the case of the two-component transcriptional system. Specifically, solving the four design problems in Propositions 4 and 5 and Corollary 1 is equivalent to tuning the ratio of the production and decay rates of the transcription factor, the ratio of the binding and unbinding rates of the transcription factor with promoters, and the total amount of DNA.

## 5 Summary and discussion

In this work, we have explored the capabilities of a method that was recently proposed by Mélykúti et al. [13,20] to derive analytical expressions for the stationary solutions to the CME. The method relies on gluing simple state spaces together recursively at one or two states. We have introduced graph theoretical characterisations of state spaces that can be constructed by performing single-state gluing of paths, cycles or both sequentially. Moreover, we have characterised a set of stochastic biochemical reaction networks for which the state spaces can be constructed using the gluing technique finitely many times. For these reaction networks, we have developed recursive algorithms that apply the gluing technique to obtain stationary distributions in analytic form. Combining recursion and asymptotic analysis, we have extended the method to derive the stationary distribution for an infinite state space, illustrating with the example of two interconnected transcriptional components. In addition, we have discussed using analytic stationary distributions to design desired distributional properties by searching over the parameter space of reaction rate constants and the amount of DNA.

While stationary distributions can be obtained symbolically by computing left null vectors of transition rate matrices, the method discussed in this work provides an alternative that is of interest for a number of reasons. Firstly, it is interesting from a theoretical perspective, as it provides a link between the graphical structure of the state space associated with the CME, and the corresponding stationary distribution. Secondly, the possibility of creating recursive algorithms that employ the repeating structure of the state space may provide better computational or analytical tractability than methods that rely on matrix algebra to compute left null vectors of transition rate matrices. Thirdly, as we have demonstrated with the example of two interconnected transcriptional components, the recursive nature of the algorithms that can be developed using this method may enable the derivation of analytic solutions for infinite state spaces.

Analytic stationary distributions of biochemical reaction systems modelled by the CME are valuable for designing user-specified distributional properties. Using analytic solutions, we can search over the parameter space of the model for stationary distributions with, for example, desired shape, modality, and moments, similar to [31]. Stationary distribution design is part of designing distributional responses of stochastic biochemical systems [32]. These methods are useful in, for instance, synthetic biology, where a principal goal is to design genetic circuits that meet user-specified design criteria. They are also important for designing population-level distributions of heterogeneous cell responses when these distributions capture unique information that is not encoded in individual cells [34].

Another application of analytic stationary distributions is to reduce the CME model using the quasi-steady-state approximation [35, 36]. This involves separating a system into fast and slow subsystems and subsequently removing the fast dynamics from the model by substituting in stationary distributions of the fast subsystems at each time step of the slow subsystem. While traditionally developed for deterministic modelling, stochastic model reduction has drawn significant attention in recent years [37], and the methods developed in this study hope to extend the set of tools for the computation of stationary distributions.

For future work, it is valuable to broaden the set of stochastic biochemical reaction networks for which analytic forms of stationary distributions can be obtained using recursive algorithms. One potential breakthrough would be to generalise Mélykúti et al.’s technique in order to allow gluing state spaces at more than two states simultaneously. Since many biochemical reactions are irreversible, it is also important to study the gluing technique on directed state spaces that are weakly connected graphs. For biochemical reaction networks with known analytic stationary distributions, a natual next step is to achieve complex distributional design by optimising over possibly large parameter spaces.

## Authors’ contributions

The project was formulated by AAB, VS, and RMM. XFM, AAB, and VS performed the mathematical analyses. XFM constructed the algorithms. AAB illustrated the design of stationary distributions. The paper was written by XFM, AAB, and VS. All of the authors edited the paper.

## Acknowledgements

The authors gratefully thank Anandh Swaminathan, Justin Bois, Enoch Yeung, Albert R. Chern, Scott C. Livingston, and Charlie Erwall for insights and comments. We thank Reza Ghaemi and Domitilla Del Vecchio for granting us permission to use the figure of two interconnected transcriptional components.

